# Urban-adapted mammal species have more known pathogens

**DOI:** 10.1101/2021.01.02.425084

**Authors:** Gregory F. Albery, Colin J. Carlson, Lily E. Cohen, Evan A. Eskew, Rory Gibb, Sadie J. Ryan, Amy R. Sweeny, Daniel J. Becker

## Abstract

The world is rapidly urbanising, inviting mounting concern that urban environments will experience increased zoonotic disease risk. Urban animals could have more frequent contact with humans, and therefore may transmit more zoonotic parasites; however, these animals have a specific set of underlying traits that may determine their parasite burdens while predisposing them to urban living, and they may be subject to more intense research effort, both of which could complicate our ability to reliably identify the role of urbanisation in driving zoonotic risk. Here, we test whether urban mammal species host more known zoonotic parasites, investigating the potential underlying drivers while accounting for a correlated suite of phenotypic, taxonomic, and geographic predictors. We found that urban-adapted mammals have more documented parasites, and more zoonotic parasites specifically: despite comprising only 157 of the 2792 investigated species (6%), urban mammals provided 39% of known host-parasite combinations and showed consistently higher viral discovery rates throughout the last century. However, contrary to predictions, much of the observed effect was attributable to research effort rather than to urban adaptation status itself, and urban-adapted species in fact hosted fewer zoonoses than expected given their total observed parasite richness. We conclude that extended historical contact with humans has had a limited impact on the number of observed zoonotic parasites in urban-adapted mammals; instead, their greater observed zoonotic richness likely reflects sampling bias arising from proximity to humans, which supports a near-universal underlying pattern of conflation between zoonotic risk, research effort, and synanthropy. These findings underscore the need to resolve the ecological mechanisms underlying links between anthropogenic change, sampling bias, and observed wildlife disease dynamics.

## Introduction

As the rate of infectious disease emergence continues to rise, it is becoming increasingly important to identify and understand the drivers of zoonotic risk in wild animals (Jones *et al*. 2008; Keesing *et al*. 2010; Morse *et al*. 2012). Humans are rapidly altering patterns of wildlife disease through a combination of climate change and land conversion, both of which are expected to drive increased spillover (i.e., interspecific transmission of parasites from animals into humans (Jones *et al*. 2008; Keesing *et al*. 2010; Loh *et al*. 2015; Hassell *et al*. 2017; Carlson *et al*. 2020a; Cohen *et al*. 2020; Gibb *et al*. 2020)). Urban environments in particular are expected to facilitate the emergence of zoonotic pathogens in wildlife (Keesing *et al*. 2010; Hassell *et al*. 2017; Becker *et al*. 2018; Murray *et al*. 2019; Werner & Nunn 2020), through a combination of impaired immune systems fed by anthropogenic resources (Becker *et al*. 2015, 2018) and greater pollution (Becker *et al*. 2020a) as well as increased proximity of wild animals to humans (Hassell *et al*. 2017; Albery & Becker 2021). This combination of factors is likely to become even more problematic in the future as the world’s population continues to rapidly grow and urbanize (Seto *et al*. 2012; Chen *et al*. 2020; Gao & O’Neill 2020).

Previous meta-analyses have uncovered elevated stressors and greater parasite burdens or parasite diversity in urban animals, with the general expectation that the urban environment weakens host immune responses (Murray *et al*. 2019; Gibb *et al*. 2020; Werner & Nunn 2020). However, these studies usually comprise relatively few examples spread across a small selection of animal species, reducing their ability to generally address the question of how urbanisation affects zoonotic disease risk. Moreover, the results of such analyses have been equivocal, with both positive, negative, and neutral effects of urban living on dimensions of wildlife disease (Murray *et al*. 2019; Gibb *et al*. 2020; Werner & Nunn 2020). Testing whether urban-adapted mammal species exhibit greater zoonotic risk in a broad-scale, pan-mammalian analysis could provide more general answers to this question, informing the design of parasite sampling regimes and efforts to mitigate zoonotic disease risk in humans.

A recent pan-mammalian study used a literature review to build a database of mammal species’ urban adaptation status (i.e., their ability to live off urban resources (Santini *et al*. 2019)), which they then linked with species-level phenotypic traits. Although different traits were important for different mammalian orders, species with larger litters were generally more likely to be urban-adapted. This relationship could explain the common observation that fast-lived host species (i.e., those that favour reproduction over survival) tend to disproportionately source zoonotic parasites (Keesing *et al*. 2010; Ostfeld *et al*. 2014; Albery & Becker 2021). Complicating matters, a given species’ observed parasite diversity depends inherently on the effort that has been directed towards examining it (Olival *et al*. 2017; Gutiérrez *et al*. 2019; Teitelbaum *et al*. 2019; Mollentze & Streicker 2020). Such research effort is heterogeneously distributed in space (Allen *et al*. 2017; Olival *et al*. 2017; Jorge & Poulin 2018) and across mammal species, particularly with regards to life history (Albery & Becker 2021) and taxonomy (Olival *et al*. 2017; Mollentze & Streicker 2020). As such, sampling bias could be important in mediating observed trends among urbanisation, life history, and zoonotic parasite diversity. In particular, urban mammal species may have more zoonoses as a proportion of their known parasite richness, because historic contact with humans has allowed more parasites to spill over into humans and be observed. Although it has been shown that human-adjacent animals have both more parasite species and more zoonoses (Gibb *et al*. 2020), it is unclear yet whether human contact has filtered them to produce disproportionately more observed zoonoses in urban species. Here, we take a macroecological approach to investigate (*i*) whether urban-affiliated mammal species have more zoonotic parasites and (*ii*) whether they harbour more zoonotic parasites than expected given their overall parasite diversity. We anticipated that species capable of adapting to urban settings would host a higher diversity of known parasites, owing to greater susceptibility and more intense sampling effort, and that a disproportionately high number of these parasites would be known to be zoonotic as a result of their greater historical contact with humans. We further expected that urban adaptation status would account for some variation in the effects of life history traits on parasite richness, implying that fast-lived species more often transmit zoonotic parasites because they are more likely to inhabit urban environments in close proximity to humans (Albery & Becker 2021).

## Results

We ran a series of generalised linear mixed models (GLMMs) that broadly supported our prediction that urban-adapted mammals would have greater parasite richness. Our first model set examined parasite richness as a response variable, revealing that urban mammals have more known parasites (Figure 1A, SI1), and more zoonoses specifically (Figure 1B, SI2). This urban bias diminished substantially in magnitude when we added citation counts as an explanatory variable representing research effort (Figure 1C); in the case of overall parasite richness, adding citation counts rendered the effect of urban adaptation non-significant (P=0.07). Citation number was strongly positively associated with urban status, overall parasite richness, and overall zoonotic richness (Figure 1C, 2), as well as being significant for all parasite subgroups (Figure SI4-5). We elaborated on these models by accounting for spatial patterns in parasite richness and sampling effort using a centroid-based SPDE effect. These effects improved model fit substantially (ΔDIC>150), and increased the magnitude and significance of the urban adaptation effects (Figure 1C; P=0.018 and 0.006). As such, we conclude that urban species have slightly higher parasite diversities when sampling effort and geographic heterogeneity are accounted for.

**Figure 1:**
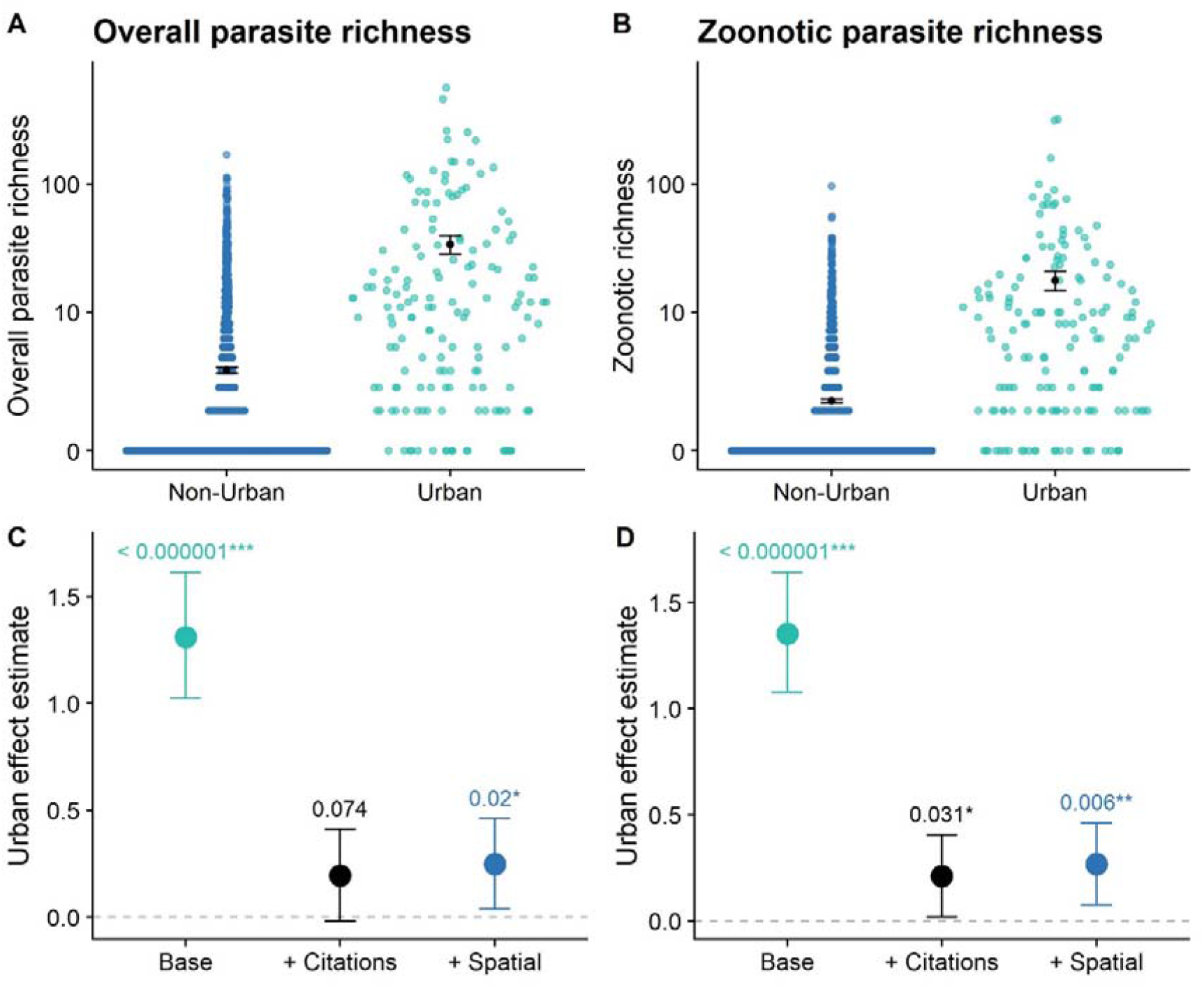
Urban-adapted mammals have more known parasites (A, C) and zoonoses specifically (B, D). In A-B, each point represents a mammal species, stratified by species that can capitalize on urban environments and those that do not. The Y axis represents the species’ known parasite diversity, on a log10 scale. Black dots and error bars represent raw group means and standard errors, respectively. Panels C-D present the urban adaptation effect for overall richness (C) and zoonotic richness (D), across multiple model formulations. The “base” models include all fixed effects but citation number; the following model includes citation number; and the third includes both citation number and a spatially distributed SPDE random effect. Points represent the mean of the posterior effect estimate distribution from the GLMMs; error bars represent the 95% credibility intervals. Numbers above the error bars display the P values, with asterisks denoting levels of significance (*<0.05; **<0.01; ***<0.001).

**Figure 2:**
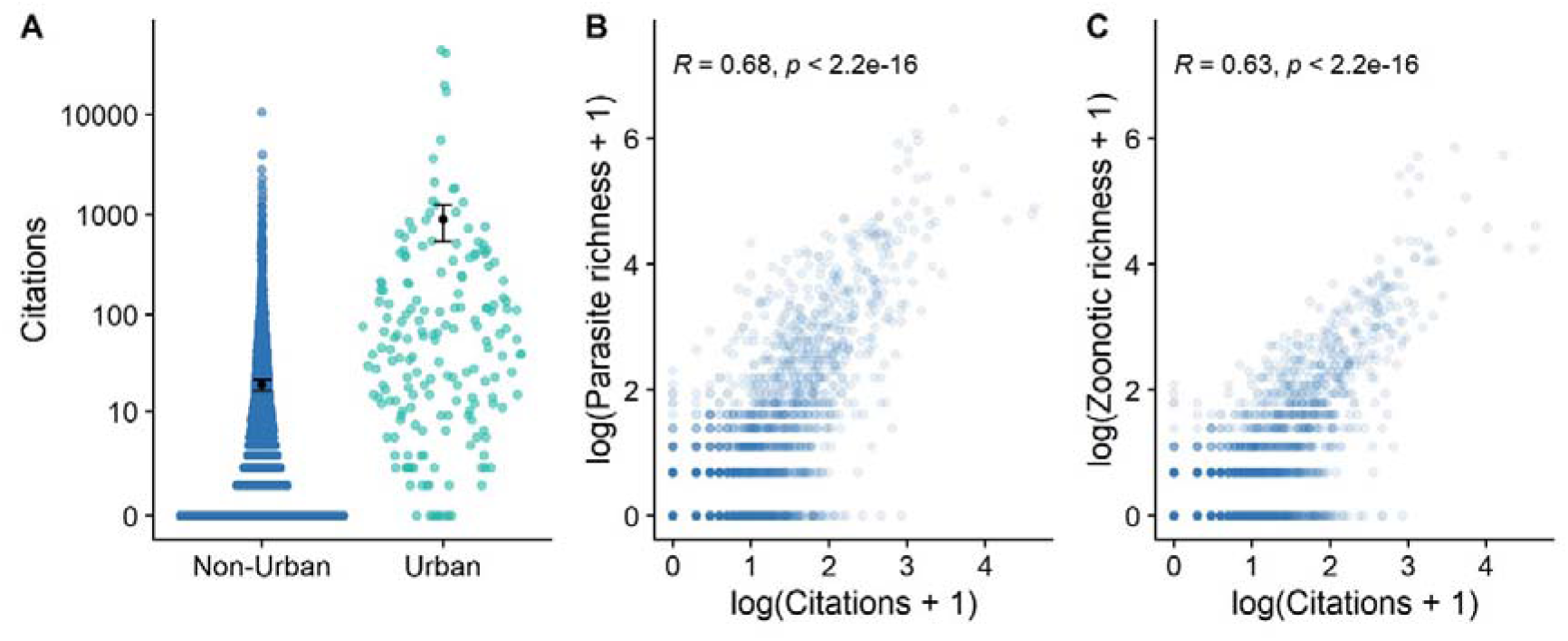
Citation numbers are higher in urban species (A), and drive observed parasite richness (B) and observed zoonotic parasite richness (C). Each point represents a species. R and P values are derived according to Spearman’s rank correlations. In panel A, black dots and error bars represent raw group means and standard errors, respectively. See Figure 1 for the slope estimates from the GLMMs for panels B-C.

To provide further insight into how histories of sampling may have shaped current patterns of observed pathogen richness across urban-adapted and non-urban species, we used our dataset to descriptively visualise historical pathogen discovery rates and publication effort trends (1930-2015), following a recent study of mammalian viral discovery (Gibb et al. 2021). We find that fewer annual discoveries generally occur in urban species; however, because there are so few urban-adapted species (157 out of 2792), these species have been, on average, more intensely studied and with a higher parasite richness since the mid-1960s (Figure SI7). Notably, differences in mean parasite richness between urban-adapted and non-urban species have continued to widen in the intervening years as the discrepancy in sampling effort has continued to grow (Figure SI7). This finding suggests that higher observed parasite richness in urban-adapted species is largely driven by long-term, accumulated differences in sampling effort.

We constructed a path analysis, which showed that urban adaptation was not associated with greater zoonotic richness when accounting for a direct effect of parasite richness; in fact, the estimated effect was slightly negative (Figure 3; P=0.024). In contrast, the indirect effect of urban adaptation on zoonotic diversity acting through parasite diversity was positive, substantial, and significant (effect +0.401; 95% credibility interval 0.116-0.749; P=0.004; Figure 3). Taken together, these results imply that positive effects of urban adaptation on zoonotic diversity act largely through greater overall known parasite diversity, rather than by disproportionately elevating zoonotic parasite richness specifically. We performed multiple further analyses to examine several dimensions of urban adaptation and sampling bias that could affect our results. There was no improvement in model fit when urban status interacted with host order, suggesting that the effect of urban adaptation on parasite diversity and zoonotic risk did not vary between mammal orders (ΔDIC<5 relative to the base model). We built a generalised additive mixed model (GAMM) to next examine whether citation numbers had different effects for urban and non-urban species, but found no support for the interaction (ΔDIC<5). Similarly, multivariate models revealed concordance between estimates for the effect of urban adaptation across parasite subtypes and implied that the urban effects were not being driven by specific groups of parasites. Finally, we used zero-inflated GLMMs to account for mammal species with no recorded parasites, demonstrating strong urban biases for the count component (i.e., the number of parasites a mammal species hosted) as well as the inflation component (i.e., whether the mammal species had greater than zero known parasites; Figure SI6). This finding implies that our results are not being disproportionately driven by excess zeroes produced by the inclusion of pseudoabsences (i.e., species without any evidence of parasites).

**Figure 3:**
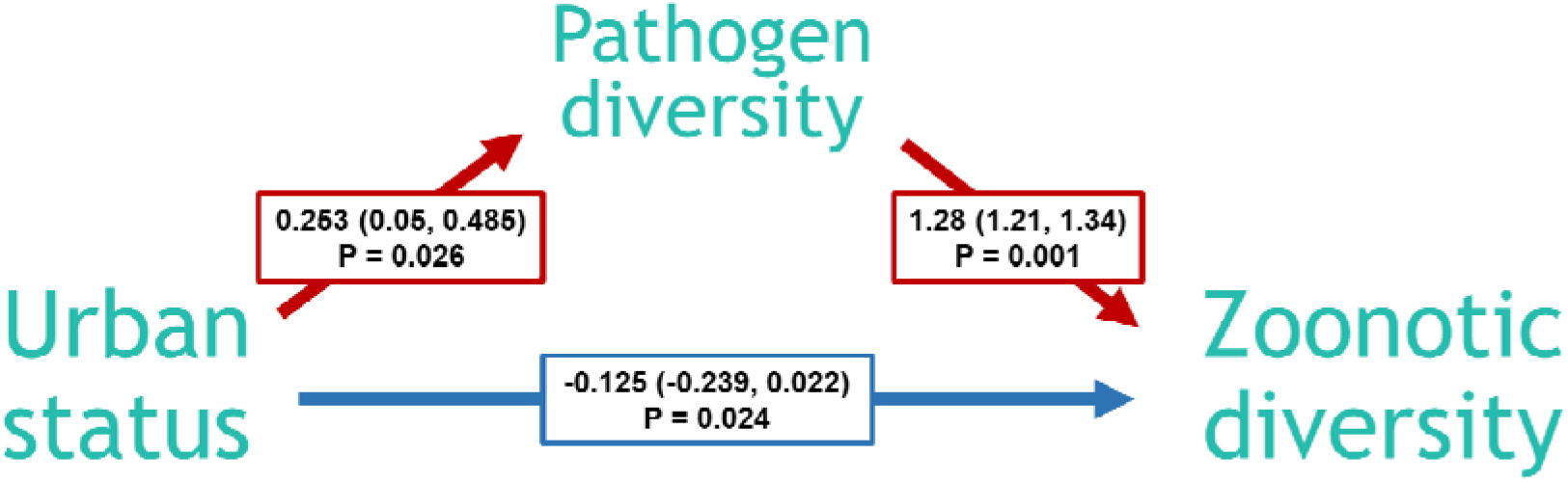
Path analysis revealed that urban-adapted mammals do not have more zoonoses than expected given their overall parasite diversity. Arrows denote hypothesised causal relationships. Red lines represent positive effects and blue lines represent negative effects. Other variables were included in the component linear models, but are not displayed in this figure for clarity. Labels display the model effect estimates on the log link scale, with 95% credibility intervals in brackets, and P values based on proportional overlap with 0.

A GLMM with different spatial fields for urban and non-urban species was not an improvement over the overall SPDE model (ΔDIC=14.35 relative to the SPDE model). This implies that the bias towards greater parasite richness in urban species is relatively evenly distributed across the globe, rather than being focussed in certain areas. These findings imply that our results were robust to geographic variation in parasite richness, and revealed strong spatial patterns (Figure 4C). We also found a substantial positive estimate for the fixed effect of absolute latitude, revealing greater known parasite diversities in temperate regions (Figure 4B). We also observed substantial between-continent variation in parasite diversity (Figure 4B): North America was associated with the greatest parasite diversity, followed by Africa, then Eurasia, South America, and Oceania.

**Figure 4:**
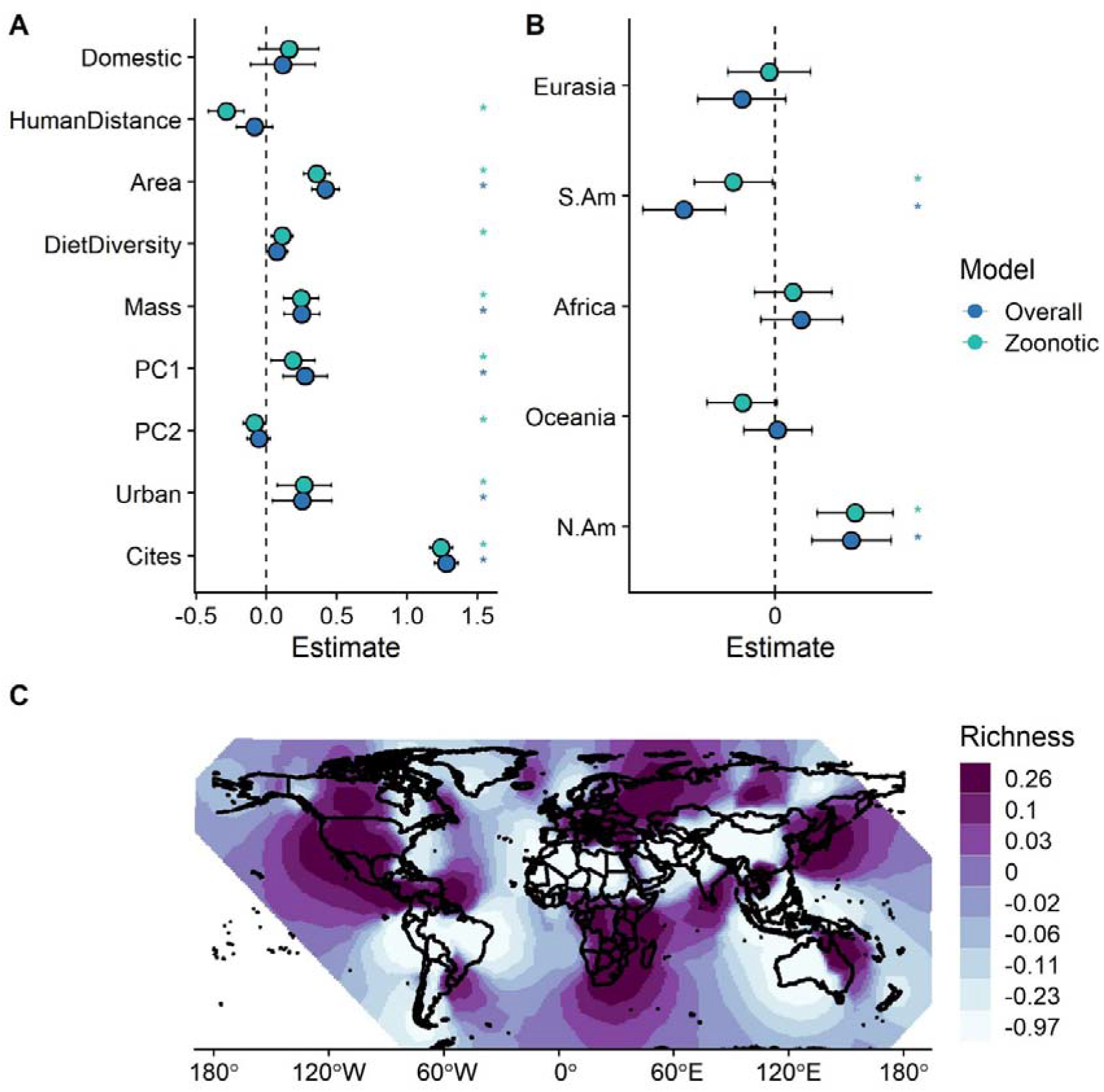
Model fixed effect estimates and spatial effects. (A) Fixed effects from the GLMMs for overall parasite richness and zoonotic richness, excluding order-level effects. These models included an SPDE random effect to control for spatial autocorrelation. (B) Fixed effect estimates from the non-spatial GLMMs for overall parasite richness and zoonotic richness. In A-B, points represent the mean of the posterior effect estimate distribution from the GLMMs; error bars represent the 95% credibility intervals. Asterisks denote estimates that were significantly different from zero. Order-level effects have been left out for clarity; see the Figures SI4-5 for full model effect estimates. (C) Spatial distribution of the SPDE random effect, identifying hot-and coldspots of parasite richness when non-spatial fixed effects (all effects except latitude and continent) are taken into account. Darker colours correspond to greater parasite richness.

Lastly, we also uncovered support for a range of other important species traits driving parasite richness (Figure 4A). Most notably, faster life history was associated with greater (zoonotic) parasite diversity, according to PC1 (Figure 4A). However, in the path analysis model, the effect of life history on zoonotic richness was supplanted by the inclusion of overall parasite richness (Figure SI3). This finding reveals that, as with urban adaptation status, life history is associated with greater overall parasite richness rather than zoonotic richness specifically. There was substantial between-order variation in zoonotic and overall diversity (Figure SI4-5), but adding a continuous phylogenetic similarity effect did not improve on the order-level effects (ΔDIC<5). Diet diversity was positively associated with zoonotic richness, but not overall parasite richness (Figure 4A). Phylogenetic distance from humans was negatively associated with zoonotic richness overall (Figure 4A), with zoonotic richness of viruses and helminths, and with overall richness of viruses and helminths; however, phylogenetic distance from humans was positively associated with overall richness of arthropods (Figure SI4-5). Greater range area was associated with increased (zoonotic) parasite richness overall (Figure 4A) and for many parasite subsets (Figure SI4-5). Finally, domesticated species had more zoonotic helminths and protozoa (Figure SI5) but did not differ in overall parasite richness from non-domesticated mammal species (Figure 4A, SI4).

## Discussion

Using a global pan-mammalian dataset of host species’ traits and parasite associations, we found that urban-adapted mammal species have more known parasites, and in turn more zoonotic parasites, arising largely from research effort. This finding builds on recent work showing that wild animals with at least one known zoonotic parasite tend to inhabit human-managed landscapes (Gibb *et al*. 2020), but we used a much broader dataset of urban-adapted mammals and applied a strict definition of urban adaptation based on long-term resource use and fitness in urban landscapes (Santini *et al*. 2019), while accounting for a correlated suite of phenotypic traits, research effort, and geographic biases, including range size and phylogenetic relatedness to humans. Additionally, we were surprised to find that urban mammals’ zoonotic richness was in fact lower than expected given their observed parasite richness. Our findings therefore do not support our main prediction that urban-adapted species host more known zoonotic parasites because they have had more historical contact with humans, creating more opportunities for the spillover of potentially-zoonotic parasites (Albery & Becker 2021). Rather, urban species appear to have been preferentially sampled for non-zoonotic parasites, likely as a result of their proximity to humans and ease of sampling – that is, mammals in urban contexts might be more often spontaneously examined for parasites, while mammals in non-urban contexts are more likely to be examined specifically when they are suspected sources of zoonotic parasites. The reason for urban mammals’ greater overall parasite richness remains uncertain, and many questions still linger about the drivers of zoonotic diversity in urban wildlife. Most pressingly, why has human-wildlife contact not driven greater zoonotic diversity in urban species?

Sampling bias is one of few universal phenomena in ecological research (Estes *et al*. 2018; Hughes *et al*. 2020), and understanding these biases is integral to designing interventions and predicting the consequences of global change. Our models revealed that urban-adapted species have been more thoroughly sampled for parasites than non-urban species, but in roughly similar patterns. Known urban status is highly geographically heterogeneous (Santini *et al*. 2019) and in a similar pattern to disease surveillance (Allen *et al*. 2017; Olival *et al*. 2017; Jorge & Poulin 2018), which we expected to be driving our perceived urban adaptation effect. The spatial patterns of parasite richness that we discovered mirror previously reported biases towards temperate, high-income countries (Titley *et al*. 2017; Hughes *et al*. 2020), and were particularly high in North America, while being particularly low in South America, confirming that parasite biodiversity is substantially undersampled in the tropics (Jorge & Poulin 2018). This reflects the pattern of urban mammal diversity, which peaks at high latitudes and is low in South America, Southeast Asia, and sub-Saharan Africa (Santini *et al*. 2019). However, accounting for this heterogeneity in fact increased the urban bias estimate rather than decreasing it. Further, there was no significant interaction of urban adaptation with either the spatial effect or host order, implying minimal geographic and taxonomic bias in these urban-directed sampling processes. Finally, our temporal analysis revealed that urban and non-urban mammals have been subjected to similar trends in parasite discovery rate over the last century, with citation counts and parasite diversity following similar shapes throughout. The only analysis that implied a qualitatively different sampling trend in urban-adapted mammal species was our path analysis, which revealed that urban-adapted species have fewer known zoonotic parasites than expected given their observed parasite richness. Taken together, the evidence suggests that urban species are much better-sampled for parasites than non-urban species, but with a stronger focus on non-zoonotic parasites, and this urban bias should be considered in future species-level analyses of zoonotic risk.

Even accounting for these layers of bias, our data still retained a positive effect of urban status, suggesting that either 1) urban mammals are subject to a specific sampling bias that could not be detected through our analyses, or 2) urban environments increase overall parasite diversity through effects on host immunity, behaviour, and demography. Although these effects did not disproportionately increase zoonotic parasite diversity, urban mammals nevertheless host many zoonotic parasites as a result of their greater overall parasite richness, and therefore understanding this trend may be important for public health. Anthropogenic pollutants, altered nutrition, and greater host densities in urban environments have been shown to weaken host immune systems and promote greater burdens and diversities of parasites when comparing hosts along urban-rural gradients (Becker *et al*. 2018; Murray *et al*. 2019). Such intraspecific effects should accordingly scale up such that urban-adapted species have greater parasite richness than species that do not experience such immune impairments. Similarly, greater host densities and resource concentrations could facilitate elevated rates of density-dependent parasite transmission within and between species, rendering urban-affiliated species more likely to maintain parasites and resulting in greater observed parasite diversity (Lloyd-Smith *et al*. 2005). However, there is some evidence that urban wildlife might exhibit stronger immunological resistance (Hwang *et al*. 2018; Strandin *et al*. 2018; Cummings *et al*. 2020), which would be expected to have the opposite effect on parasite diversity, and a previous study found that some parasite groups are decreased in urban environments rather than increased (Werner & Nunn 2020). Unfortunately, the field is generally lacking in large-scale cross-species analyses of immune function that would be required to differentiate these possibilities (Albery & Becker 2021; but see Downs *et al*. 2020a, b). Ideally, future analyses incorporating life history, habitat preference, immunity, and parasite diversity may be better able to differentiate the mechanisms underlying these species’ zoonotic risk (Albery & Becker 2021).

Achieving broad insights into the urban drivers of zoonotic risk may require finer-scale data than we had access to here. This study was conducted with a minimum compatibility filter: we considered a species as a host of a given parasite if it was observed with said parasite at any point in the literature, and richness was calculated as the sum of these associations across parasite subgroups. While studies of parasite diversity are common in macroecology, this deliberately narrow scope limits inference about a range of relevant processes including host competence (i.e., species’ ability to transmit parasites; Becker *et al*. 2020b), prevalence of the parasite in the host populations, host density, and, therefore, the *rate* of spillover (i.e., the number of animal-to-human transmission events per unit of time). These are all important components of a species’ zoonotic risk, and some hosts undoubtedly present substantial zoonotic risk despite having relatively low known parasite diversity. For example, prairie dogs (*Cynomys ludovicianus*) only have five known parasites in our dataset, yet they are a widespread and abundant species and may play an important role in epizootic outbreaks of plague (*Yersinia pestis*) in North America (Hanson *et al*. 2007). Given this disparity, it remains unclear how closely a species’ zoonotic diversity should correlate with the rate of spillover from these species; as such, we caution that our analysis does not necessarily offer insights into the relative frequency or rate of spillover events, or the potential severity of zoonotic outbreaks, in urban environments.

Providing a general answer to the question “does urbanisation increase the risk of zoonotic disease” may require datasets of individual-or population-level infection status, using multiple hosts and parasites, distributed across a wide range of urbanisation gradients. Higher-resolution datasets such as these would facilitate untangling of within-and between-species confounders, as well as accounting for spatiotemporal covariates like urban habitat composition (Gecchele *et al*. 2020). These data are increasingly publicly available and are being used in large-scale analyses of disease dynamics (e.g. (Cohen *et al*. 2020; Albery *et al*. 2021)); as such, these analyses may become increasingly possible in coming years. Regardless, in these and other analyses, correlated changes in the magnitude and shape of sampling biases (e.g. towards zoonotic versus non-zoonotic parasites) should be taken into account when examining links among anthropogenic change, wildlife disease, and zoonotic risk.

## Methods

### Data sources

#### Phylogeographic data

We used the PanTHERIA dataset (Jones *et al*. 2009) as a backbone for mammal taxonomy and phenotypic traits such as body mass. Phylogenetic data were derived from a mammalian supertree (Fritz *et al*. 2009), as used for several host-virus ecology studies (e.g. Olival *et al*. 2017; Albery *et al*. 2020; Becker *et al*. 2020). The tree’s phylogenetic distances between species were scaled between 0 and 1. Geographic data were taken from the IUCN species ranges (IUCN 2019). For each species, we calculated total range area by adding together the areas for the 25 km raster cells in which they were present.

To derive a measure of study effort, which often explains substantial variation in parasite diversity (Olival *et al*. 2017; Mollentze & Streicker 2020), we conducted systematic PubMed searches to identify how many publications mentioned a given mammal species, following previous methodology (Becker *et al*. 2020b). Domestication status used a *sensu lato* definition based on whether a species has ever been partially domesticated, coded as a binary variable. For example, despite being widespread in the wild, the European red deer (*Cervus elaphus*) is coded as “Domestic” because it is often farmed, notably in New Zealand (Mason 1994). Because we were investigating spatial distributions of species (see above), fully domesticated species that do not exist in the wild (e.g. cattle, *Bos taurus*) were generally excluded due to their absence from the IUCN species ranges. To investigate whether dietary flexibility could affect parasite diversity, following previous methodology (Santini *et al*. 2019), we derived diet diversity by calculating a Shannon index from the EltonTraits database proportional diet contents (Wilman *et al*. 2014).

#### Life history data

To investigate how host life history variation affects parasite richness, we used a previously published, mass-corrected principal components analysis (PCA) of life history variation across mammal species (Plourde *et al*. 2017). The first two principal components (PCs) from this analysis, which explained 86% of variation in six life history traits (Plourde *et al*. 2017), were used as explanatory variables in our models. The six life history traits were gestation length, litter size, neonate body mass, interbirth interval, weaning age, and sexual maturity age. PC1 explains 63% of the variance in the six traits, representing a generalisable slow-fast life history axis. PC2 explains 23% of variance in these traits and represents greater investment in gestation time and larger offspring. Both PCs were available for all mammals in our dataset. We coded the PCs such that increasing values corresponded to “faster” life history (i.e., favouring greater reproduction over survival).

#### Urban adaptation data

We identified each species’ habitat preferences using a published database of long-term urban adaptation status in mammals (Santini *et al*. 2019). This dataset was compiled using literature searches to identify species that were observed inhabiting urban environments; species are either coded as a “visitor” or a “dweller”, based on whether they rely fully on urban environments to survive and reproduce (dweller) or whether they continue to rely on non-urban resources (visitor). This approach distinguishes our analysis from previous studies (e.g. Gibb et al., 2020): we use a strict definition of “urban-adapted” species, defining them as “mammals that survive, reproduce, and thrive in urban environments,” rather than basing urban status purely on survey records collected in urban settings. All species that were in PanTHERIA but were not in the urban adaptation dataset were coded as “non-urban”. We used urban adaptation as a binary variable, coding species as 0 or 1 depending on whether it was in the urban adaptation dataset. Overall, 180 species in our dataset were coded as a 1, denoting that they had been observed living off urban resources.

#### Host-parasite association data

The recently released CLOVER dataset (Gibb *et al*. 2021) is the most comprehensive open-source dataset on the mammal-virus network. Here, we use an expanded version of this dataset that encompasses all parasites, rather than restricting to viruses, making our analysis the first analytical study to use these taxonomically broad parasite data. This dataset was synthesized from four large-scale datasets of host-parasite associations, each collected through a combination of web scrapes and systematic literature searches (Wardeh *et al*. 2015; Olival *et al*. 2017; Stephens *et al*. 2017; Shaw *et al*. 2020). These include the Enhanced Infectious Diseases Database (EID2; Wardeh *et al*. 2015); the Host-Pathogen Phylogeny Project (HP3; Olival *et al*. 2017); the Global Mammal Parasite Database (GMPD; Stephens *et al*. 2017); and a large-scale database on viruses and bacteria and their known hosts (Shaw *et al*. 2020). These contain a range of parasite groups, including viruses, bacteria, protozoa, fungi, helminths, and arthropods. In this conjoined dataset, host-parasite associations were counted according to demonstrated compatibility: that is, if a host species had ever been discovered infected with a given parasite, it was coded as a 1, and all undemonstrated associations were assumed absent. In addition to the taxonomic reconciliation underlying the CLOVER dataset, we cleaned the parasite names with the R package *taxize* (Chamberlain & Szöcs 2013), removing parasites that were not identified to species level and ensuring that no parasites existed under multiple identities. This ensured that no host-parasite associations were counted twice, resulting in a total 18,967 unique host-parasite associations.

From this conjoined dataset, we derived the following traits for each mammal host species in our dataset: 1) **Total parasite richness**: the number of unique parasite species known to infect a given host species; 2) **Zoonotic parasite richness:** the number of these parasites that has also been observed to infect humans in our dataset. All analyses were repeated for overall parasite numbers (e.g., total number of zoonoses across all parasite groups) and for specific parasite subgroups (viruses, bacteria, protozoa, fungi, helminths, and arthropods).

For each analysis, to facilitate model fitting, we eliminated species for which there were missing data and then removed all host orders for which there were fewer than 20 species or for which fewer than 1% of species had one or more known parasites. Leaving these taxa in did not notably alter fixed effects estimates generally but generated unlikely estimates for order-level effects). When combining the phenotypic, urban adaptation, and parasite datasets, any species with no known parasite associations was coded as a zero (i.e., a pseudoabsence), under the assumption that species with no known parasites are still informative of variables associated with low parasite richness (Albery & Becker 2021).

## Models

### Base model

To analyse associations between urban adaptation status and parasite richness, we used Generalised Linear Mixed Models (GLMMs) inferred using Integrated Nested Laplace Approximation (INLA) (Lindgren *et al*. 2011; Lindgren & Rue 2015). We used two response variables with a negative binomial distribution: total parasite richness and zoonotic parasite richness, where the second value was a subset of the first. Explanatory variables included: Citation number (log(x+1)-transformed); Host order (7 levels: Artiodactyla, Carnivora, Chiroptera, Lagomorpha, Primates, Rodentia, Soricomorpha); Urban adaptation status (binary; non-urban/urban); range area (continuous, log-transformed, defined above); Phylogenetic distance from humans (continuous, scaled 0-1); Body mass (continuous, log-transformed); Domestication status (binary); and two life history principal components (PC1 and PC2; continuous, taken from Plourde et al. 2017). We also applied these models to each parasite subset to assess the generality of our parameter estimates. To examine how much of the observed urban effects were attributable to research effort, we

### Urban:citation GAMs

Because urban status and citation number were highly correlated and showed very different distributions, we fitted a generalised additive model (GAM) that was otherwise identical to our GLMMs, but with a smoothed term for citations that included an interaction with urban status.

### Urban-order interaction model

We then compared the base model with one including an interaction between host order and urban adaptation status to investigate whether the effect of urban adaptation varied taxonomically. We used the Deviance Information Criterion (DIC) to measure model fit, with a threshold change (ΔDIC) under 5 denoting competitive models.

### Phylogenetic model

For each model, we fitted a phylogenetic similarity effect in place of the host order effect to estimate how phylogenetic relatedness between species contributed to similarity in parasite richness. We used DIC to identify whether this effect improved model fit in the same way as the interaction model.

### Multivariate models

To investigate whether urban adaptation status had different effects for the richness of different parasite types, we fitted two multi-response models using the *MCMCglmm* package (Hadfield 2010): one for overall richness and one for zoonotic richness. These models used each of the six parasite groups as response variables and included the same fixed effects, with different (but correlated) slopes for each response. Comparing the model’s estimates for the effect of urban adaptation for each parasite allowed us to ask whether specific parasite groups are significantly more likely to be associated with urban adaptation status than others.

### Zero-inflated models

To investigate whether pseudoabsences were disproportionately altering our results, we ran zero-inflated models of parasite and zoonotic richness again using *MCMCglmm* to control for processes that specifically generate zero-counts. These models generate two estimates for each explanatory variable: 1) the effect on the probability that a species’ parasite count is greater than zero (“zero-inflation”) and 2) the effect on parasite count greater than zero when accounting for this effect (“Poisson”). Importantly, the Poisson component of this model generates some zeroes itself, which improves upon similar models (e.g. hurdle models) in which all zeroes must be produced by the inflation term. This model allows us to identify whether, for example, urban species are simply more likely to have one or more known parasites, rather than having a greater overall known parasite richness, and whether our choice to code mammals with no known parasites as zero-counts would influence the results.

### Historical rates of parasite discovery

To investigate how differences between urban and non-urban wild mammals have accumulated over time, we analysed historical rates of parasite discovery and citation effort (from PubMed) between 1930 and 2020, following the methodology described in Gibb et al. 2021. Briefly, each unique host-parasite association was assigned a “discovery date” (the year of the earliest reported association in our dataset, based on either publication year, accession year or sampling year depending on the original data source; see Gibb *et al*. (2021) for details). We accessed yearly counts of citations from the PubMed database per host species using the `rentrez` package (Winter 2017). We visualised annual trends in novel parasite discovery and novel host-parasite association discovery in both urban and non-urban mammal species. We then fitted generalised additive models with a nonlinear effect of year (specified as a penalised thin-plate regression spline) to estimate the annual species-level mean publications, cumulative publications, parasites discovered and cumulative parasite richness, fitting separate models for urban-adapted (n=146) and non-urban (n=1365) species in our host-parasite dataset. We visualised fitted trends in these metrics to examine how differences in yearly and cumulative publication effort and parasite discovery rates have varied between urban and non-urban species (Figure SI7).

### Path analysis

To investigate whether urban mammals had a disproportionately high zoonotic richness when accounting for overall parasite richness, we fitted a path analysis (Shipley 2009) with zoonotic richness as the ultimate response variable, log(overall richness +1) as an explanatory variable, and every other explanatory variable described above. We took 1000 random draws from the posterior distributions of 1) the effect of urban affiliate status on overall parasite diversity; 2) the effect of urban affiliate status on zoonotic richness; and 3) the effect of overall richness on zoonotic richness. This approach allowed us to identify whether urban adaptation had a significant positive effect on zoonotic richness when accounting for its effect on parasite richness as a whole, informing us as to whether a disproportionate number of urban mammals’ known parasites are known zoonoses.

### Spatial model

Observed parasite diversity in mammals is highly spatially heterogeneous at a global level (Allen *et al*. 2017; Olival *et al*. 2017; Carlson *et al*. 2020b), while the diversity of known urban-adapted species is heavily biased towards North America and Eurasia (Santini *et al*. 2019). Both are driven by a combination of geographic variation in sampling effort as well as biotic and abiotic factors. To control for these spatial heterogeneities, we fitted spatial explanatory variables using three approaches. First, we 1) used a stochastic partial differential equation (SPDE) effect in INLA (Lindgren *et al*. 2011; Lindgren & Rue 2015). This effect used species’ geographic centroids in their IUCN ranges to control for spatial autocorrelation in the response variable according to Matern correlation, where species that were closer in space would be predicted to have similar numbers of known parasites as a result of sampling bias and biological factors. We first fitted one spatial field to the whole dataset to look for overall spatial structuring, and we then allowed this spatial effect to vary for urban and non-urban species to investigate whether the distribution of known richness varies between these hosts. We also 2) incorporated species’ presence on each of five continents (Eurasia, Africa, North America, South America, and Oceania) as binary variables and 3) added absolute latitude (i.e. distance from the equator). For the latter two approaches, we also fitted an interaction with urban adaptation to investigate whether the effect of urban adaptation status varied across space.

## Authorship Statement

GFA and DJB conceived the study, and GFA analysed the data and wrote the manuscript. All other authors offered thoughts on the analysis and commented on the manuscript.

## Data and Code Availability

The code used here is available at github.com/gfalbery/UrbanOutputters. The CLOVER dataset is available at github.com/viralemergence/clover.

## Acknowledgements

This work was supported by funding to the Viral Emergence Research Initiative (VERENA) consortium, including NSF BII 2021909.

## Supplement

**SIFigure 1:**
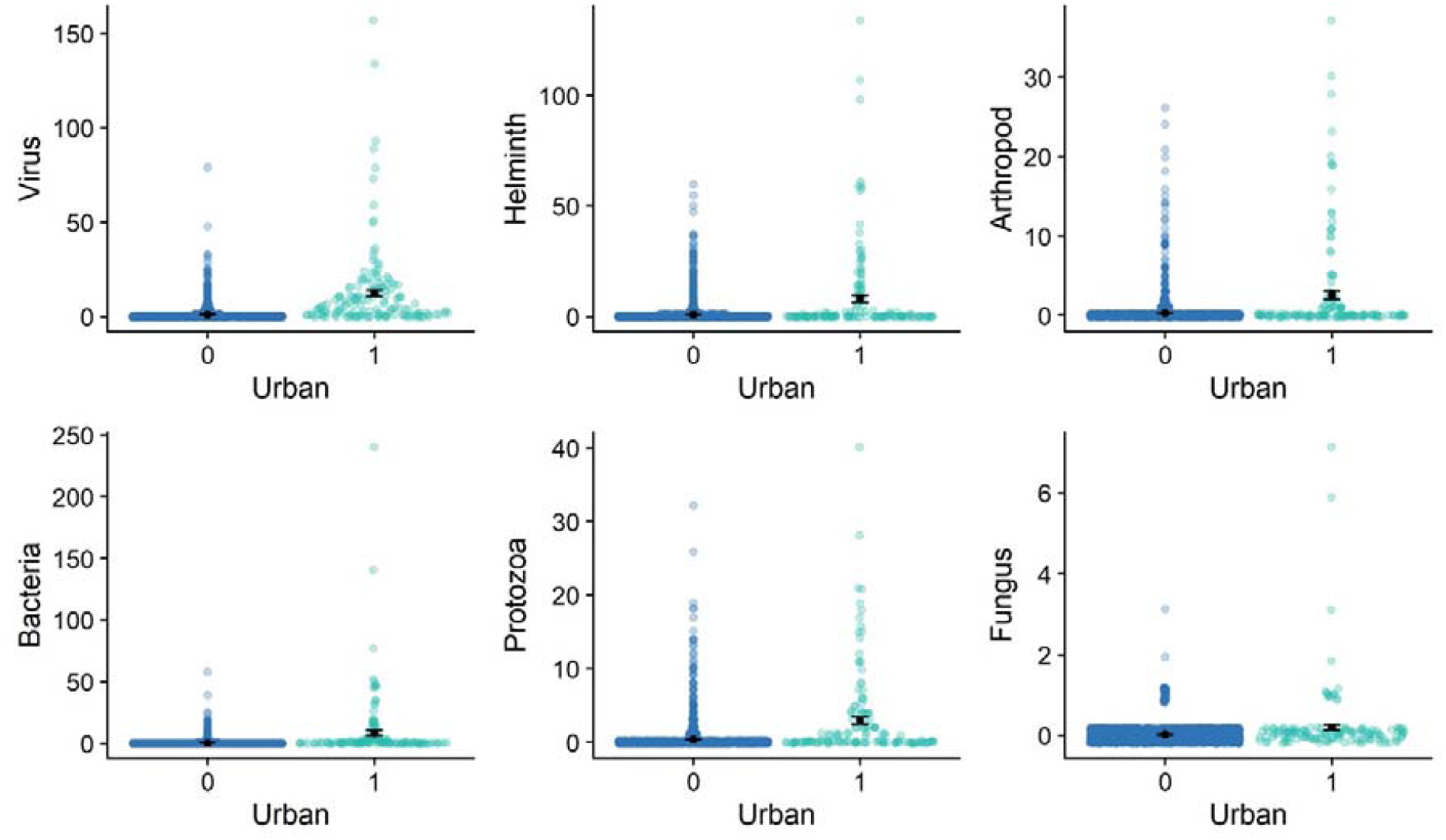
The effect of urban affiliation on diversity of pathogen subsets. Each point represents a mammal species, stratified by species that can capitalize on urban environments (1) and those that do not (0). The Y axis represents the species’ known pathogen diversity. Black dots and error bars represent raw group means and standard errors, respectively. Displayed at the top of each panel are effect sizes for the between-group difference, 95% credibility intervals (in brackets), and P values, taken from our GLMMs including other explanatory variables.

**SIFigure 2:**
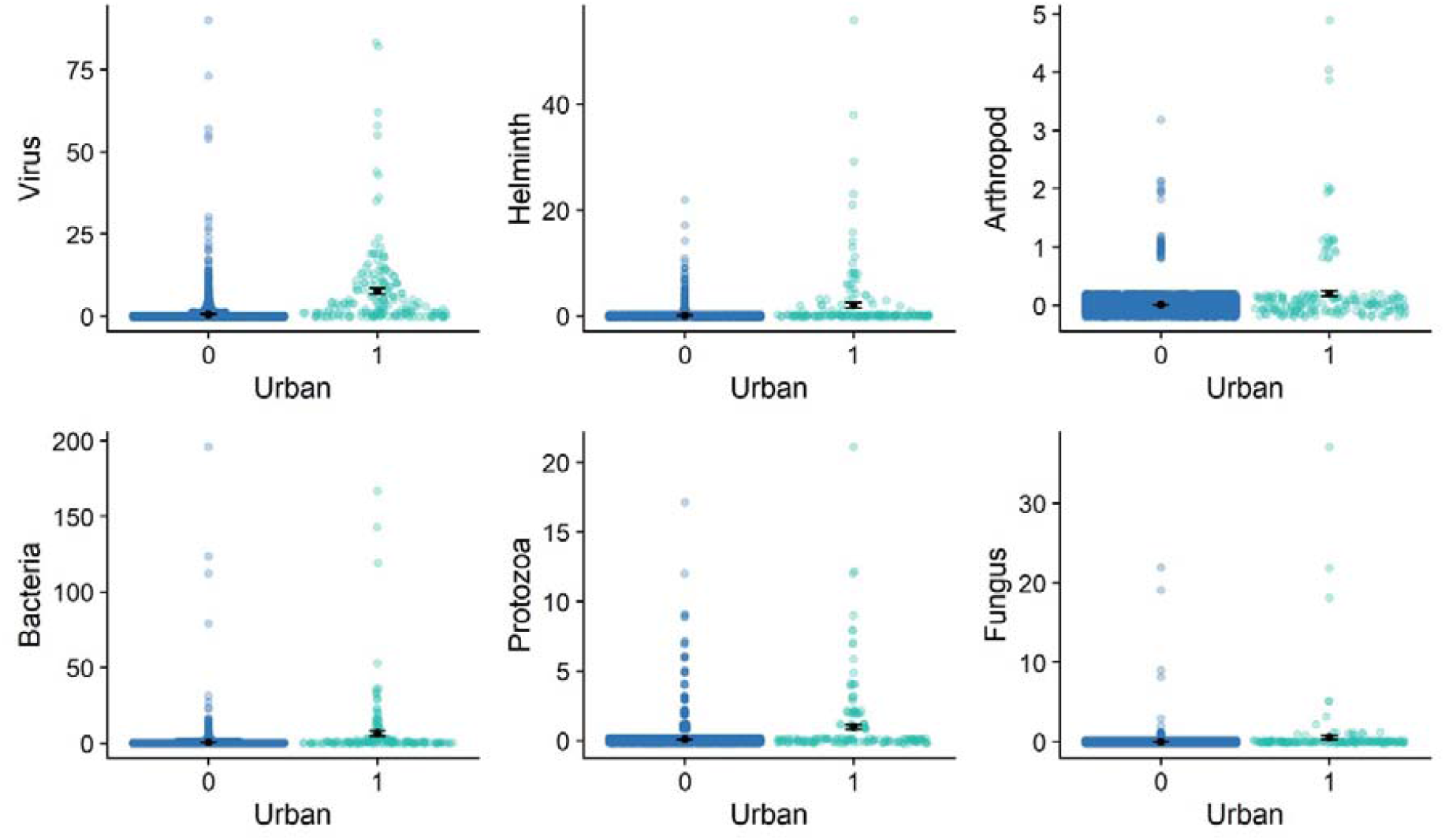
The effect of urban affiliation on zoonotic diversity of pathogen subsets. Each point represents a mammal species, stratified by species that can capitalize on urban environments (1) and those that do not (0). The Y axis represents the species’ known pathogen diversity. Black dots and error bars represent raw group means and standard errors, respectively.

**SIFigure 3:**
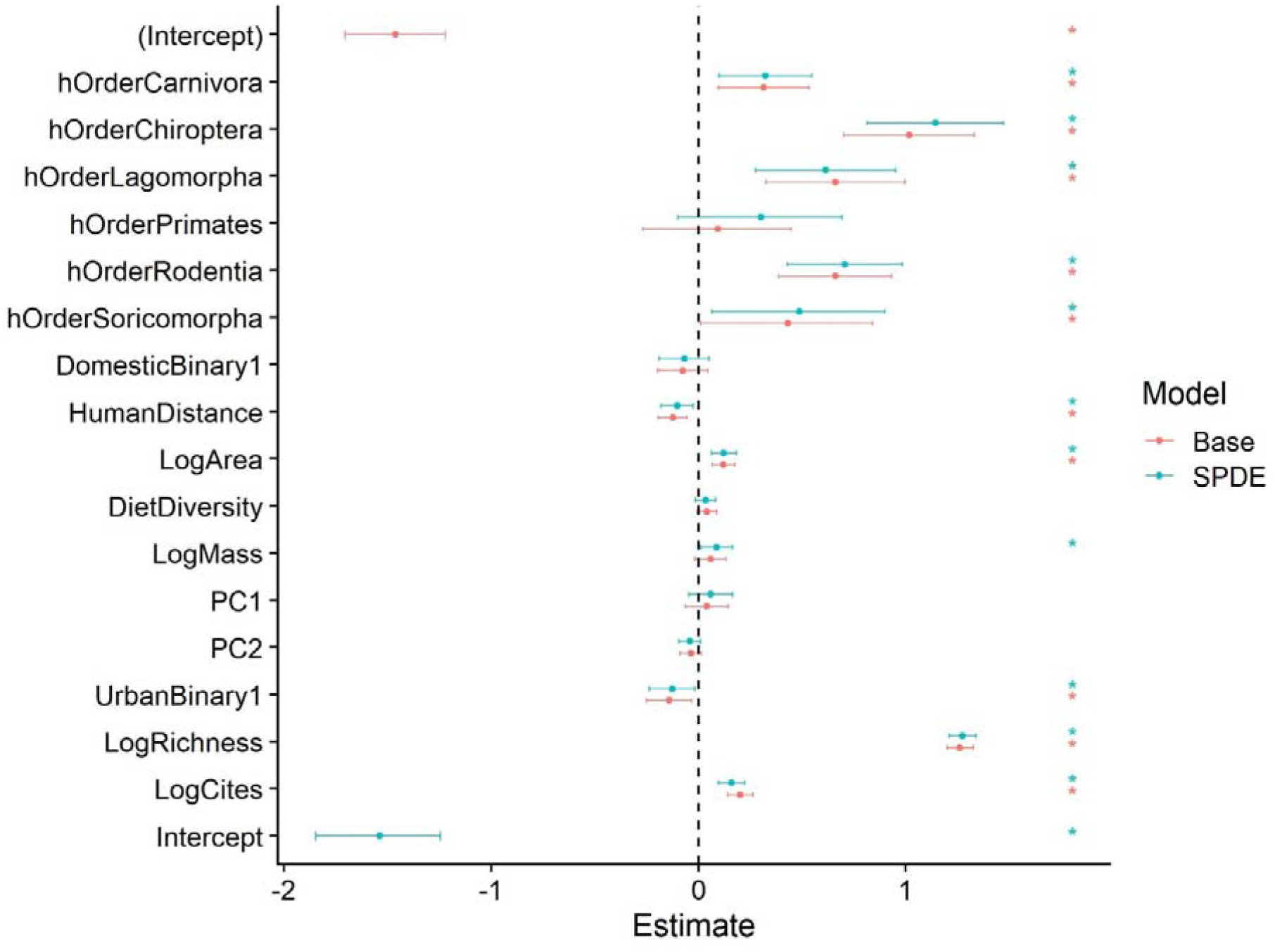
Model effects for all fixed effects retained in the path analysis models for overall zoonotic diversity, for both base and spatial model formulations. Points represent the mean of the posterior effect estimate distribution; error bars represent the 95% credibility intervals. Model effects are displayed on the link scale. Explanatory variables are described in the methods. hOrder = host order; LogCites = log(citation number + 1); DomesticBinary1 = domestic species; HumanDistance = phylogenetic distance from humans; LogArea = log(area of IUCN range) in KM^2^; DietDiversity = diet diversity; LogMass = log(body mass) in kg; PC1 = first principal component of life history traits PCA; PC2 = second principal component of life history traits PCA; UrbanBinary1 = Urban adapted species; LogRichness = log(overall parasite richness + 1).

**SIFigure 4:**
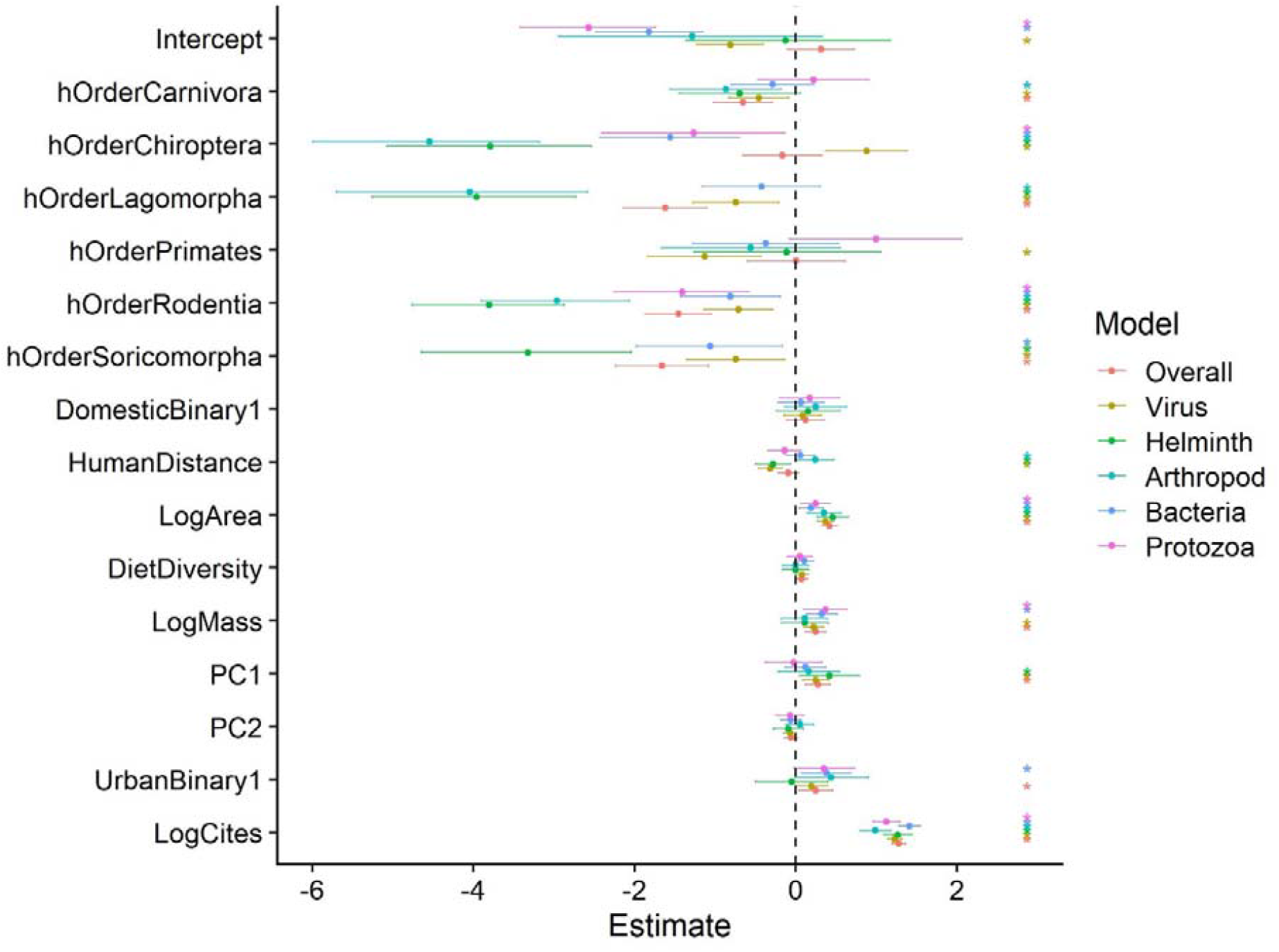
Model effects for all fixed effects in the spatial models of parasite diversity. Points represent the mean of the posterior effect estimate distribution; error bars represent the 95% credibility intervals. Different colours represent different parasite groups, including overall parasites and a range of subgroups. Model effects are displayed on the link scale. Explanatory variables are described in the methods. hOrder = host order; LogCites = log(citation number + 1); DomesticBinary1 = domestic species; HumanDistance = phylogenetic distance from humans; LogArea = log(area of IUCN range) in KM^2^; DietDiversity = diet diversity; LogMass = log(body mass) in kg; PC1 = first principal component of life history traits PCA; PC2 = second principal component of life history traits PCA; UrbanBinary1 = Urban adapted species.

**SIFigure 5:**
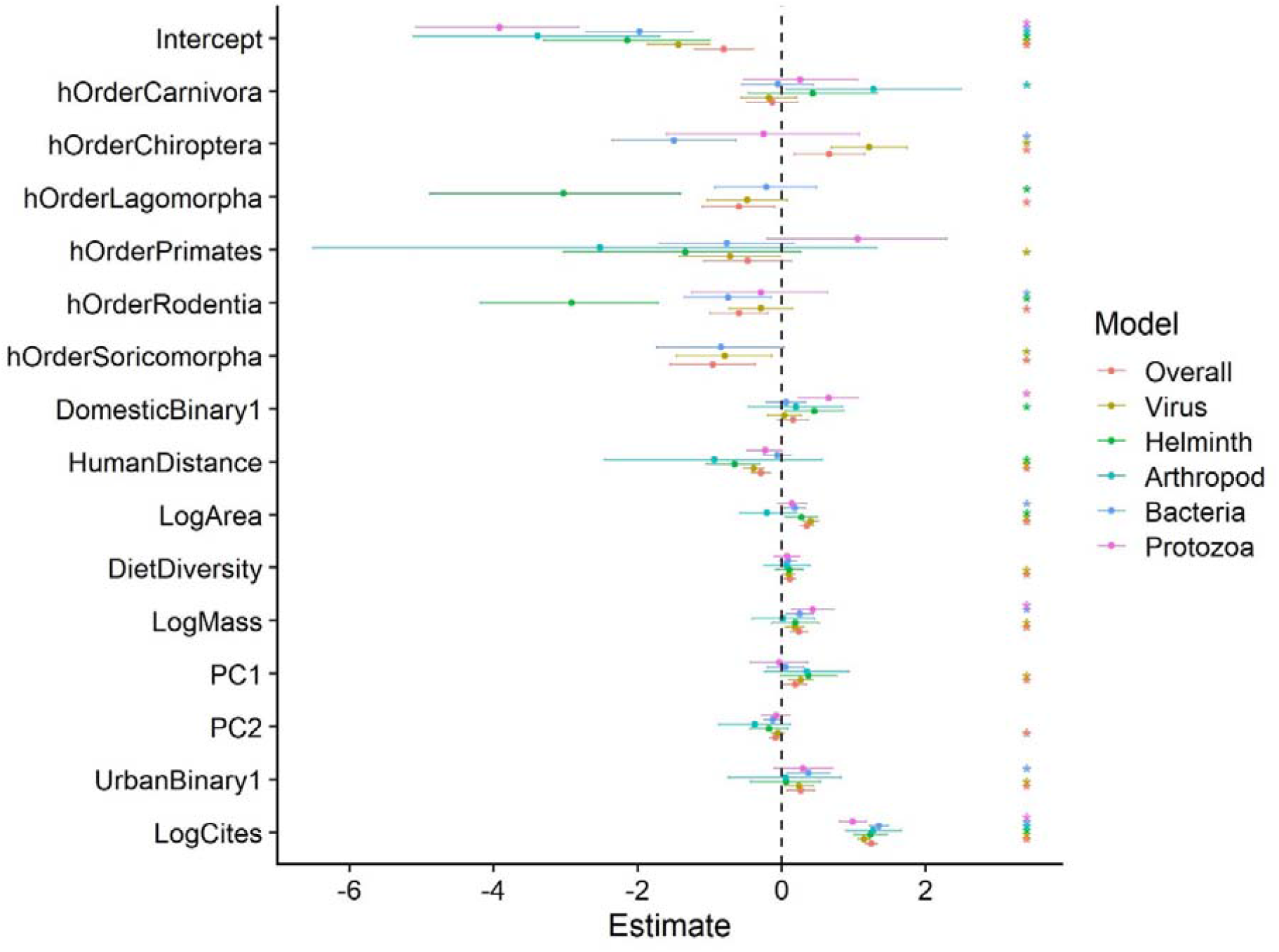
Model effects for all fixed effects in the spatial models of zoonotic parasite diversity. Points represent the mean of the posterior effect estimate distribution; error bars represent the 95% credibility intervals. Different colours represent different parasite groups, including overall parasites and a range of subgroups. Model effects are displayed on the link scale. Explanatory variables are described in the methods. hOrder = host order; LogCites = log(citation number + 1); DomesticBinary1 = domestic species; HumanDistance = phylogenetic distance from humans; LogArea = log(area of IUCN range) in KM^2^; DietDiversity = diet diversity; LogMass = log(body mass) in kg; PC1 = first principal component of life history traits PCA; PC2 = second principal component of life history traits PCA; UrbanBinary1 = Urban adapted species.

**SIFigure 6:**
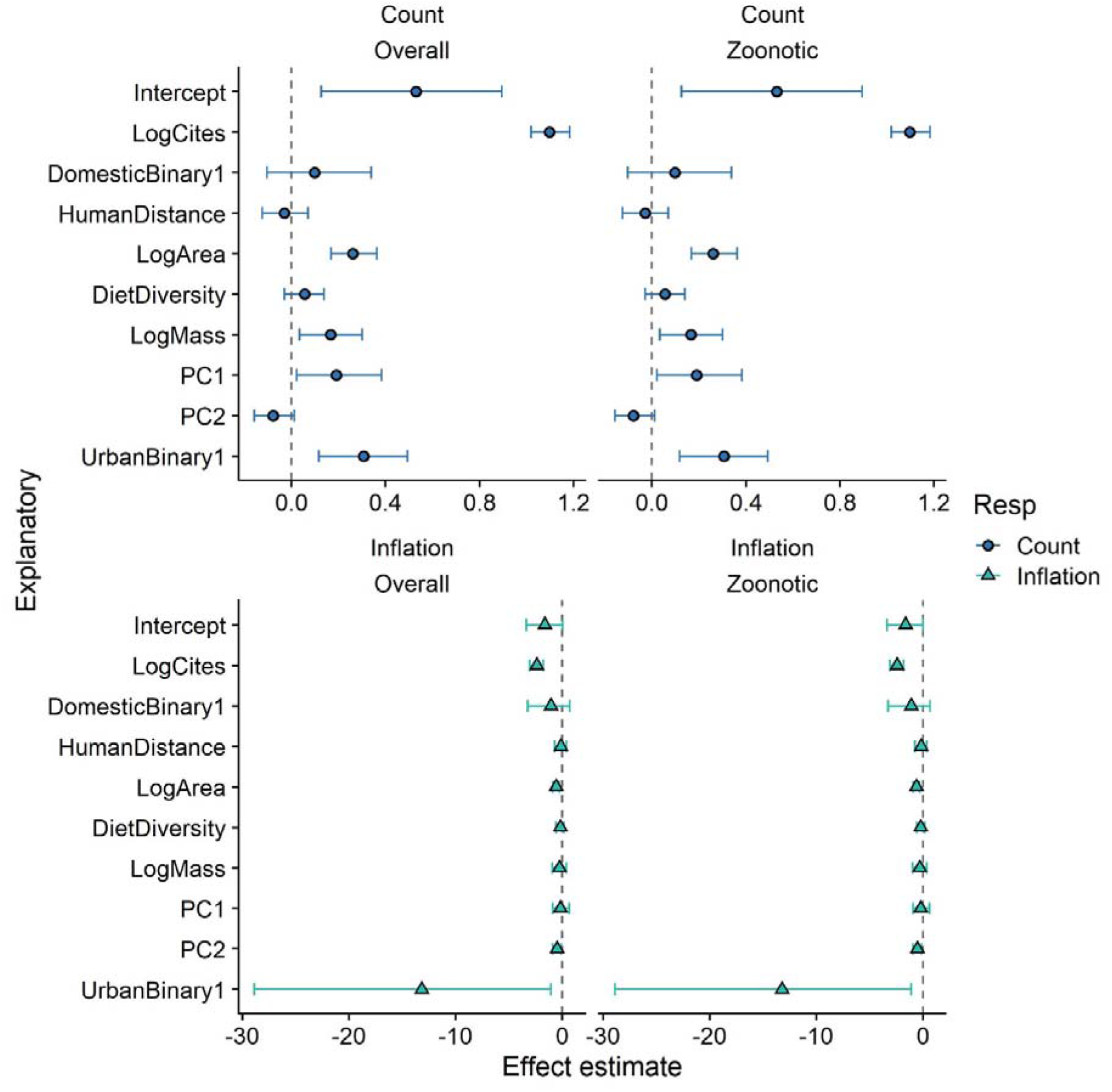
Model effects for all fixed effects in the zero-inflated models for overall parasite diversity (left) and zoonotic parasite diversity (right), for both the count components (top) and the zero-inflation component (bottom). Points represent the mean of the posterior effect estimate distribution; error bars represent the 95% credibility intervals. NB the inflation estimates represent the probability that a given species has zero known parasites, so can be interpreted as the inverse of the count estimates. Model effects are displayed on the link scale. Explanatory variables are described in the methods. LogCites = log(citation number + 1); DomesticBinary1 = domestic species; HumanDistance = phylogenetic distance from humans; LogArea = log(area of IUCN range) in KM^2^; DietDiversity = diet diversity; LogMass = log(body mass) in kg; PC1 = first principal component of life history traits PCA; PC2 = second principal component of life history traits PCA; UrbanBinary1 = Urban adapted species.

**SIFigure 7:**
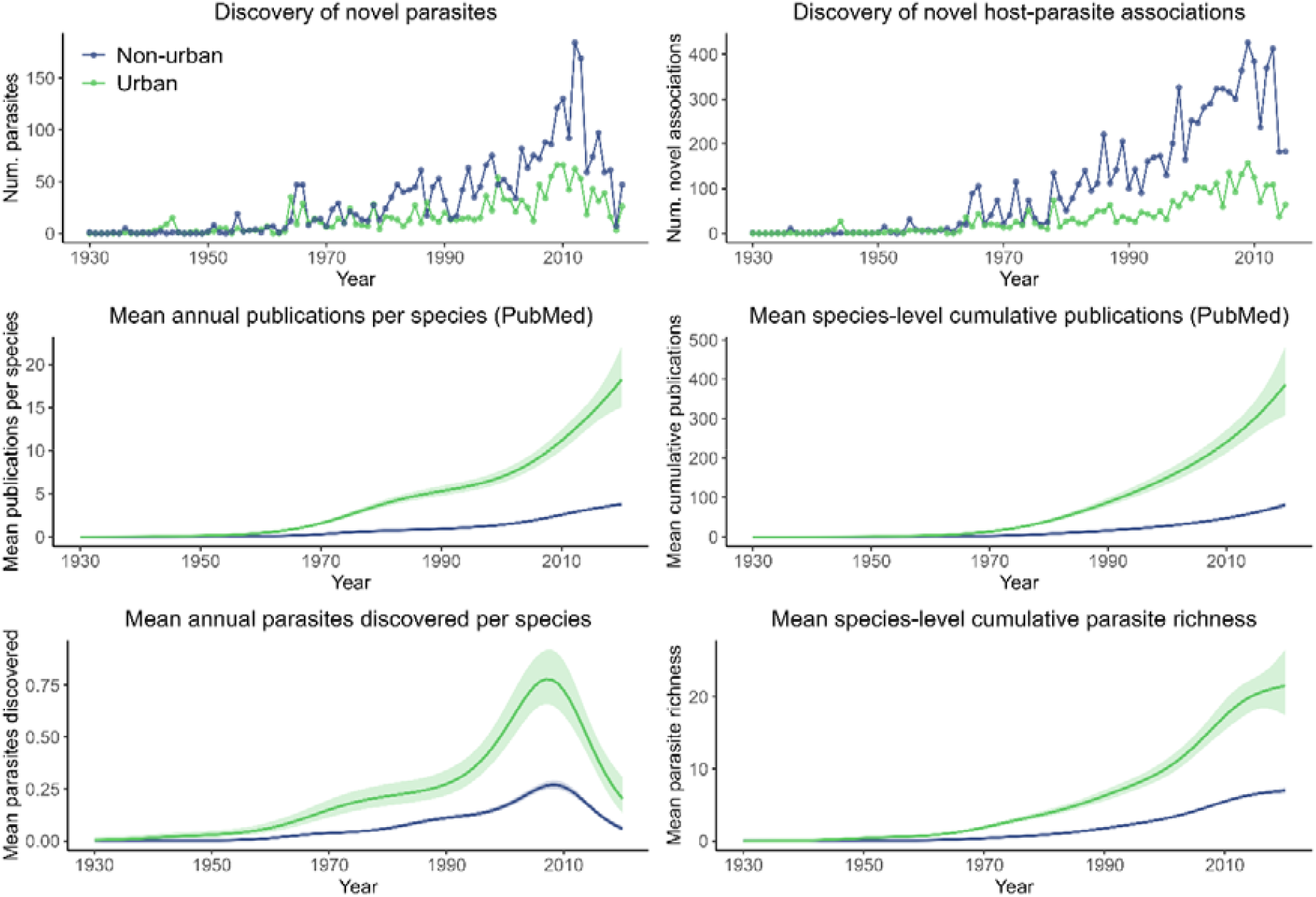
Historical trends in parasite discovery and publication effort across urban-adapted and non-urban mammals. Top row shows the annual number of novel parasites (left) discovered in either non-urban (blue) or urban (green) mammal cohorts, with a novel discovery defined as the first time a particular parasite was discovered in any species within that group, and the annual number of novel host-parasite associations (right) discovered among urban and non-urban mammals. The middle row shows modelled trends in mean species-level annual PubMed-derived publication counts (left panel) and cumulative publications (right panel) across all urban (n=146) and non-urban host species, estimated via generalised additive models with a nonlinear effect of year (see Methods). The bottom row shows modelled trends in mean species level parasite discovery (parasites per year; left panel) and cumulative parasite richness (right panel) across all urban and non-urban host species.

